# The stage- and sex-specific transcriptome of the human parasite *Schistosoma mansoni*

**DOI:** 10.1101/2023.05.02.539039

**Authors:** Sarah K. Buddenborg, Zhigang Lu, Geetha Sankaranarayan, Stephen R. Doyle, Matthew Berriman

**Author notes:** corresponding authors: Sarah Buddenborg & Matthew Berriman.

## Abstract

The flatworm *Schistosoma mansoni* is an important but neglected pathogen that causes the disease schistosomiasis in millions of people worldwide. The parasite has a complex life cycle, undergoing sexual reproduction in a mammalian host and asexual replication in a snail host. Understanding the molecular mechanisms the parasite uses to transition between hosts and develop into dimorphic reproductively competent adults may reveal new strategies for control. We present the first comprehensive transcriptomic analysis of *S. mansoni*, from eggs to juvenile worms. Focusing on eight life stages spanning free-living water-borne and parasitic stages from both intermediate and definitive hosts, we have generated deep RNA-seq data for five replicates per group for a total of 75 data sets. The data were produced using a single approach to increase the accuracy of stage-to-stage comparisons and made accessible via a user-friendly tool to visualise and explore gene expression (https://lifecycle.schisto.xyz/). These data are valuable for understanding the biology and sex-specific development of schistosomes and the interpretation of complementary genomic and functional genetics studies.

## Background & Summary

Schistosomiasis is a neglected tropical disease caused by parasitic flatworms of the genus *Schistosoma* (Phylum: Platyhelminthes, Class: Trematoda). The disease is primarily controlled via large-scale preventative chemotherapy programmes that distribute the anthelmintic drug praziquantel and in 2021, over 75 million people in 51 countries received treatment^1^. *Schistosoma mansoni* is the most widespread and most studied schistosome species, which infects humans and causes intestinal schistosomiasis, characterised by abdominal pain and diarrhoea leading to liver enlargement, hypertension of the abdominal blood vessels, and spleen enlargement in chronic cases.

Schistosomes have a complex life cycle. Sexually mature, paired adult parasites excrete eggs in the faeces of their mammalian host that, upon water contact, hatch to release motile larvae called miracidia. The miracidia seek and infect freshwater intermediate host snails and undergo asexual clonal replication before re-entering the water as cercariae to infect the mammalian definitive host. Within the mammalian host, cercariae develop into schistosomula and traverse the vasculature until juvenile male and female parasites pair and mature, becoming reproductively competent in the mesenteric veins of the intestines (**Fig. 1**).

**Figure 1:**
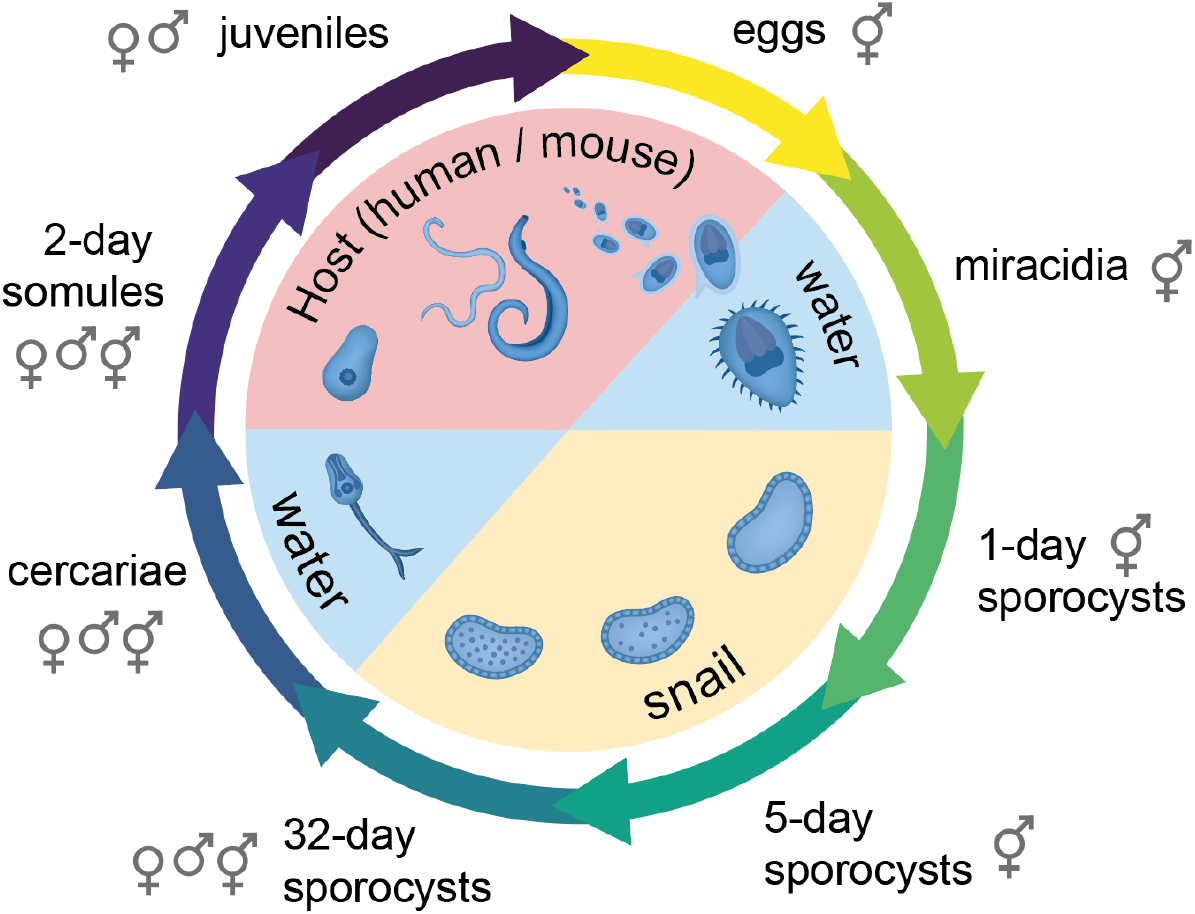
Sampling key stages of the *Schistosoma mansoni* life cycle. The flatworm *S. mansoni* undergoes a series of morphological transitions throughout development, using waterborne stages to transition between the intermediate snail host and the definitive mammalian host. Here, the life stages and sex (male, female, mixed) are shown for samples collected and sequenced using RNA-seq. Figure adapted from Laura Olivares Boldú / Wellcome Connecting Science.

Genomics approaches have provided great insight into the evolutionary history and life history traits of *Schistosoma* spp. For *S. mansoni*, this has been supported by ongoing efforts to develop high-quality reference genome resources^2–4^. As these genome resources have evolved, several studies have focused on characterising the transcriptome of various stages throughout the life cycle^4–10^, highlighting key genes involved in developmental transitions from free-living to parasitic stages and commitment towards sexual maturity. More recently, single-cell RNA sequencing on mammalian-infective stages^11–13^ and miracidia^14^ has been demonstrated, providing intricate details of body plans and cellular functions. However, a fundamental limitation of these existing transcriptomic data is that there is significant biological and technical variation across datasets, including but not limited to different parasite strains, numbers of replicates, and sequencing technologies and sampling depth, making it challenging to understand transcriptional changes across the entire life cycle or to compare data sets. Furthermore, most data sets have been sampled from pools of male and female parasites, limiting understanding of sex-specific transcriptomic changes across the life cycle.

Here, we present a comprehensive sex- and stage-specific RNA-seq data set to uncover transcriptional changes throughout the life cycle of *S. mansoni*. Our sampling (n=75) of *in vitro* and *in vivo* cultured parasites includes five replicates each of mixed-sex eggs, miracidia, primary sporocysts, secondary sporocysts, cercariae, and somules, as well as separate female and male secondary sporocysts, cercariae, somules, and liver-stage juvenile worms (sexually naïve). To aid data exploration and reuse, we have implemented a simple tool – https://lifecycle.schisto.xyz/ – to visualise and subset transcript counts from these deep-coverage Illumina sequencing data. The data reported here will support future studies on understanding the sex-specific development of schistosomes and aid in interpreting functional studies describing gene function, drug-target discovery, and single-cell interpretation.

## Methods

### Ethics Statement

All mouse procedures were conducted by authorised personnel under the Home Office Project Licence No. P77E8A062, held by Dr Gabriel Rinaldi (Wellcome Sanger Institute). All protocols were approved by the Animal Welfare and Ethical Review Body (AWERB) of the Wellcome Sanger Institute as required by the UK Animals (Scientific Procedures) Act 1986 Amendment Regulations 2012.

### Collection of parasite samples

The complete *S. mansoni* life cycle was maintained at the Wellcome Sanger Institute. The *S. mansoni* NMRI stain (Puerto Rican) was used to infect female CD-1 outbred mice (Charles River, Harlow, UK) between 8-12 weeks old and the *Biomphalaria glabrata* BB02 strain.

#### Eggs

Patently-infected mice at 42 days post-exposure were culled using an overdose of pentobarbitone containing 10 U/ml heparin. The livers were dissected and washed in a series of 1x PBS with 2% antibiotic-antimycotic and 70% ethanol. The mouse livers were minced using a sterile disposable scalpel, enzymatically degraded with collagenase, and passed through a 150 µm sieve. Schistosome eggs were purified using density centrifugation through Percoll^15,16^. Eggs from a pool of five livers of mice exposed to the same mixed-sex collection of cercariae represent one biological replicate.

#### Miracidia

Eggs from five livers for each replicate, prepared as above, were hatched under direct light at ∼28°C in MilliQ water in a 1 L flask^17,18,19^. Miracidia were collected every 30 minutes for three hours.

#### Primary (mother) sporocysts (1-day & 5-day; in vitro)

Miracidia were transformed *in vitro* by centrifugation and then resuspended in sporocyst culture medium, which was maintained at 28°C under malaria gas (90-92% N, 5% CO_2_, and 3-5% O_2_) with daily medium change^20–22^.

#### Secondary (daughter) sporocysts (32-day; snail)

For mixed-sex infections, individual *B. glabrata* snails were each exposed to 15 miracidia. For single-sex infections, each *B. glabrata* snail was exposed to a single miracidium. Each biological replicate represents an infection initiated from a different miracidia pool. Clonal, asexual reproduction of the parasite within the snail host results in a single-sex infection, and the cercariae developed from mono miracidium-infected snails are all single-sex. At 32 days post-exposure to miracidia/miracidium, *B. glabrata* snails were dissected, and secondary sporocysts (also known as daughter sporocysts) were collected with special attention to removing all visible snail tissue.

#### Cercariae

After ∼4-8 weeks post-exposure to miracidia/miracidium, snails begin to release cercariae into the water. Cercariae were released by exposing patent snails to bright light at 28°C for up to 2 hours.

#### Somules

Mixed-sex or single-sex cercariae were shed from snails, as above, and placed on ice for 30 mins. Cercariae tails were removed by repeated centrifugation and resuspension in 1x PBS with 2% antibiotic-antimycotic. The parasites were placed in schistosomula medium at 37°C and 5% CO_2_ overnight. The following day, the separated tails were removed with a transfer pipet. Transformed schistosomula were maintained *in vitro* at 37°C and 5% CO_2_ with daily media changes^18,23–26^.

#### Juvenile worms

Mice were percutaneously infected with mixed-sex cercariae by exposing the shaved abdomens of anaesthetised mice to 250 cercariae for 30 mins. At 26 days post-exposure, sexually naïve worms were recovered via portal perfusion with approximately 30 ml of Dulbecco’s Modified Eagle Medium (DMEM; high glucose) and 10 U/ml heparin^18,27^.

After collection, all stages were gently centrifuged, washed with 1x PBS, and then stored at -80°C in 500 µl of TRIzol Reagent (Invitrogen) until RNA extraction.

### Discrimination of sex of early monomorphic life stages

Early parasite life stages are sexually monomorphic and cannot be differentiated morphologically until the juvenile adult stages. To identify both male and female stages from 32-day sporocysts (after clonal amplification) onward, we determined the sex of shedding cercariae from snails^28,29^ after mono-miracidium infections using a modified HOTSHOT method (https://health.uconn.edu/mouse-genome-modification/protocols/hotshot-method-of-dna-preparation/) to obtain DNA followed by PCR-amplification of the female-specific W1 repeat^30–32^. Snails shedding single-sex parasites were then used to generate sex-defined parasite material for subsequent stages.

### RNA isolation, library preparation, and sequencing

In preparation for RNA isolation, samples were thawed on ice, transferred to MagNA Lyser Green Beads (Roche) tubes, and homogenised using the FastPrep-24 instrument (MP Biomedical) for 2 × 30 secs. An additional 500 µl of TRIzol was added to each sample, and the homogenate was allowed to sit at room temperature for five minutes to complete the dissociation of nucleoprotein complexes. RNA was isolated using TRIzol Reagent (Invitrogen) manufacturer’s instructions.

After isolation, RNA was immediately cleaned and concentrated using the RNA Clean and Concentrator-5 kit (Zymo Research) with on-column DNase I treatment according to the manufacturer’s instructions. Total RNA was quantified on the Qubit 3 Fluorometer (Invitrogen), and quality was analysed on the 2100 Bioanalyzer with RNA 6000 Nano-chips (Agilent Technologies). RNA was stored at -80°C until library preparation.

RNA-seq libraries were prepared with the NEBNext Ultra II Directional RNA Library Prep Kit for Illumina (New England Biolabs). Sequencing libraries were quantified by qPCR, and their concentration was normalised before equimolar pooling. Libraries were sequenced using 150 bp paired-end read chemistry on two S4 lanes of an Illumina NovaSeq 6000 platform.

### RNA-seq data analysis

Demultiplexed FASTQ reads were trimmed using TrimGalore v0.4.4 (https://github.com/FelixKrueger/TrimGalore). Trimmed sequencing reads were mapped to the *S. mansoni* reference genome v10^3^ from WormBase ParaSite (https://parasite.wormbase.org/Schistosoma_mansoni_prjea36577/Info/Index/)^33^ using STAR v2.7.9a^34^ (parameters: --alignIntronMin 10 --outSAMtype BAM SortedByCoordinate --limitBAMsortRAM 90000000000). The *S. mansoni* annotation was used to guide read mapping; the available GFF was converted to a GTF using gffread from the cufflinks (v2.2.1) package^35^. Mapped read sets belonging to the same sample were combined using samtools (v1.9) merge. Protein-coding features were quantified from the results of STAR using StringTie v2.1.4^36^.

Principal component analysis (PCA) was performed in R (version 4.1.3) using the prcomp() command and the log2-transformed transcript per million (TPM) count matrix. To generate the heatmap, pairwise sample distances (type=euclidean), using TMP values of the top 25% most variable genes, were calculated in R using the dist() package.

### RNA-seq data visualisation

We used Seurat v4.2.1 (https://satijalab.org/seurat/)^37^ to construct an object using the log2-transformed TPM count matrix (log_2_(TPM+1)). Metadata, including sex, stages, and sample identities, were added accordingly. To view the data in a lower dimensionality space, top 2000 variable features were extracted using the FindVariableFeatures() function and data were then scaled for PCA analysis. The resulting Seurat object with 2D PCA embeddings, log_2_(TPM+1) data, and metadata information was deployed using the interactive visualisation tool, Cirrocumulus (The Broad Institute, Inc. and The General Hospital Corporation; https://github.com/lilab-bcb/cirrocumulus) with customisations, and can be explored at https://lifecycle.schisto.xyz.

## Data Records

The data set consists of bulk RNA-seq data from 75 samples, representing eight life stages, each with five replicates. For egg, miracidia, and 1-day and 5-day sporocysts, samples were mixed sex. For 32-day sporocysts, cercariae, 2-day somules and 26-day juveniles, pooled male, pooled female, and mixed-sex stages were collected.

Sample metadata, including read counts, mapped reads, and ENA accession information, are described in Supplementary Table 1. All sequence data can be obtained from the European Nucleotide Archive (ENA) under project accession ERP115565^38^. The *S. mansoni* v10 genome and annotation used in the analysis are available from WormBase ParaSite release 18 (https://parasite.wormbase.org/Schistosoma_mansoni_prjea36577/Info/Index/).

## Technical Validation

The dataset consists of approximately 4.4×10^9^ sequencing reads, with an average of 58.9 million reads per sample (range: 21.3–183.3 million reads). Per sample, on average, 67.4% of reads mapped to the v10 assembly of the *S. mansoni*; for most life stages (excluding 32-day sporocysts), this percentage was higher (average: 77.1%); however, 32-day sporocysts showed a considerably lower percentage of mapped reads (average: 28.1%), which can be attributed to contamination from the snail host.

To assess the replicability of the data, we calculated TPM values per gene for each sample, which we analysed by principal component analysis (PCA; **Fig. 2A**) and clustering of Euclidean distance (**Fig. 2B**). PCA, which explained 70% of the variance in the first two principal components, showed clear clustering of replicate samples with only minor overlap between some life stages; however, sex differences within a life stage were less discernible. Differentiating stage and sex was more apparent based on the clustering of TPM by Euclidean distance, with clear discrimination of cercariae, juveniles, and eggs. Two-day somules and 32-day sporocysts were also clearly differentiated by life stage; however, each contained some misclustering based on sex (single replicate for mixed sex/female for 2-day somules; two replicates for male / female samples 32-day sporocysts), emphasising the importance of our relatively high number of biological replicates. Similarly, there was some misclustering between a single replicate of miracidia and 1-day sporocysts and a single 1-day and 5-day sporocyst; however, both cases represent transitions between adjacent stages in the life cycle. Overall, most replicates are clustered with their corresponding sex- and stage-specific data sets.

**Figure 2.**
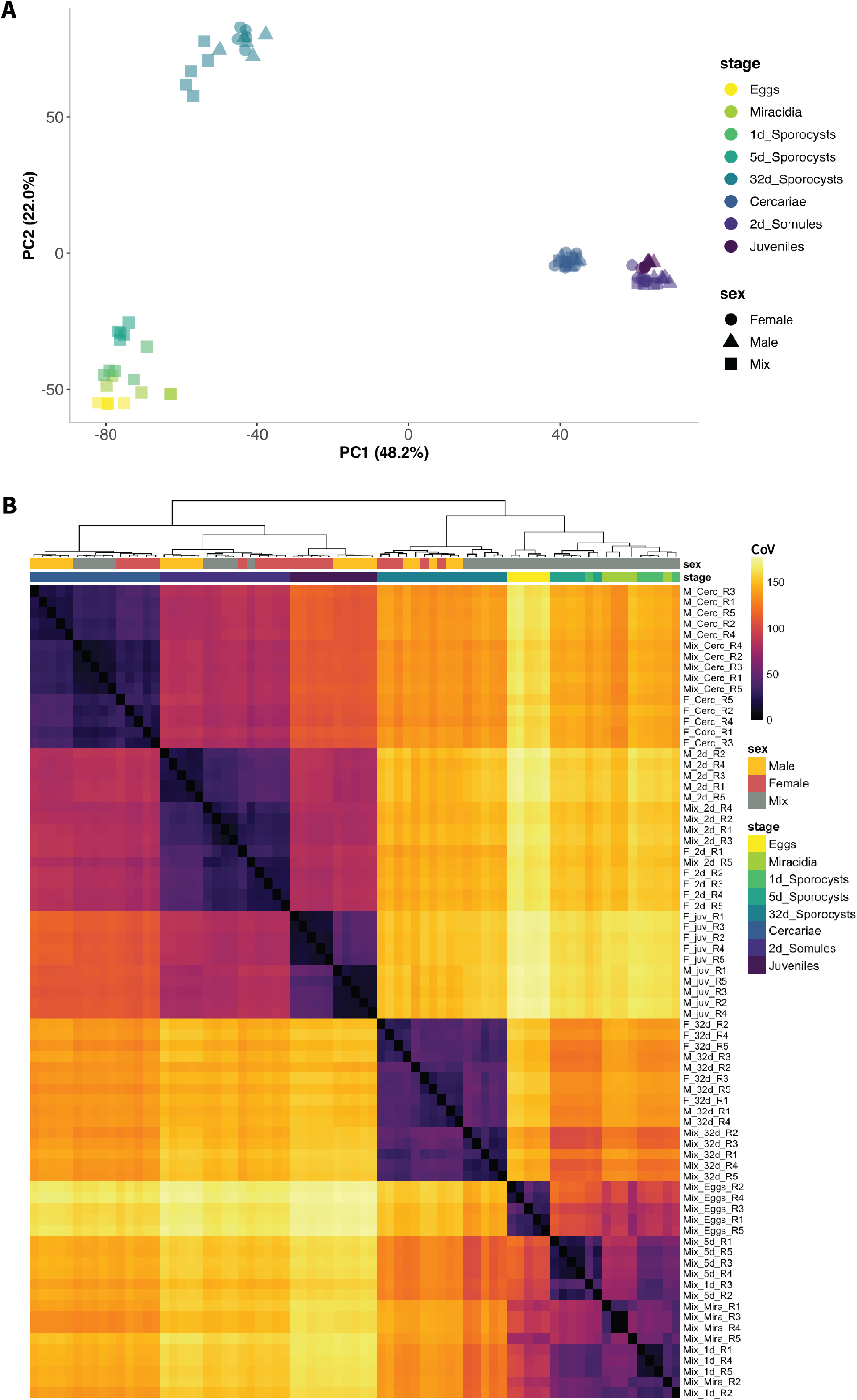
Clustering of RNA-seq data by parasite sex and life stage. **(A)** A PCA plot of the 25% most variable genes was calculated by the coefficient of variation, highlighting life stage (colour) and sex (shape). **(B)** Clustering of samples based on Euclidean distance (Coefficient of Variation; CoV) of 25% most highly variable genes. In both panels A and B, the colour is consistent with Figure 1.

## Usage Notes

These data provide a rich resource to understand the sex and stage-specific developmental changes of *S. mansoni* throughout its life cycle. To aid data exploration and reuse, we have implemented a simple tool to visualise and subset these data: https://lifecycle.schisto.xyz/. To illustrate, we highlight variation in the expression of *S. mansoni* Kunitz protease inhibitors that are proposed to be involved in the defence mechanisms of the parasite within the mammalian host^39^; in the v5 genome assembly, only a single copy was identified, whereas in the v10 genome assembly, 11 copies were identified^3^. Here, we demonstrate two approaches to visualise samples based on sex (**Fig. 3A**) and life stage (**Fig. 3B**), and differences in gene expression, either by colouring the PCA based on levels of gene expression represented as log_2_(TPM+1) (**Fig. 3C**) or by displaying mean log_2_(TPM+1) (combining each life stage) as a dotplot heatmap (**Fig. 3D**). Further customisation (colours, plots) and filtering (min, max expression, conditions) of the data are possible depending on the user’s preference to explore gene expression of any gene present in the genome. These analyses identify four groups of genes with distinct stage-specific expression patterns.

**Figure 3.**
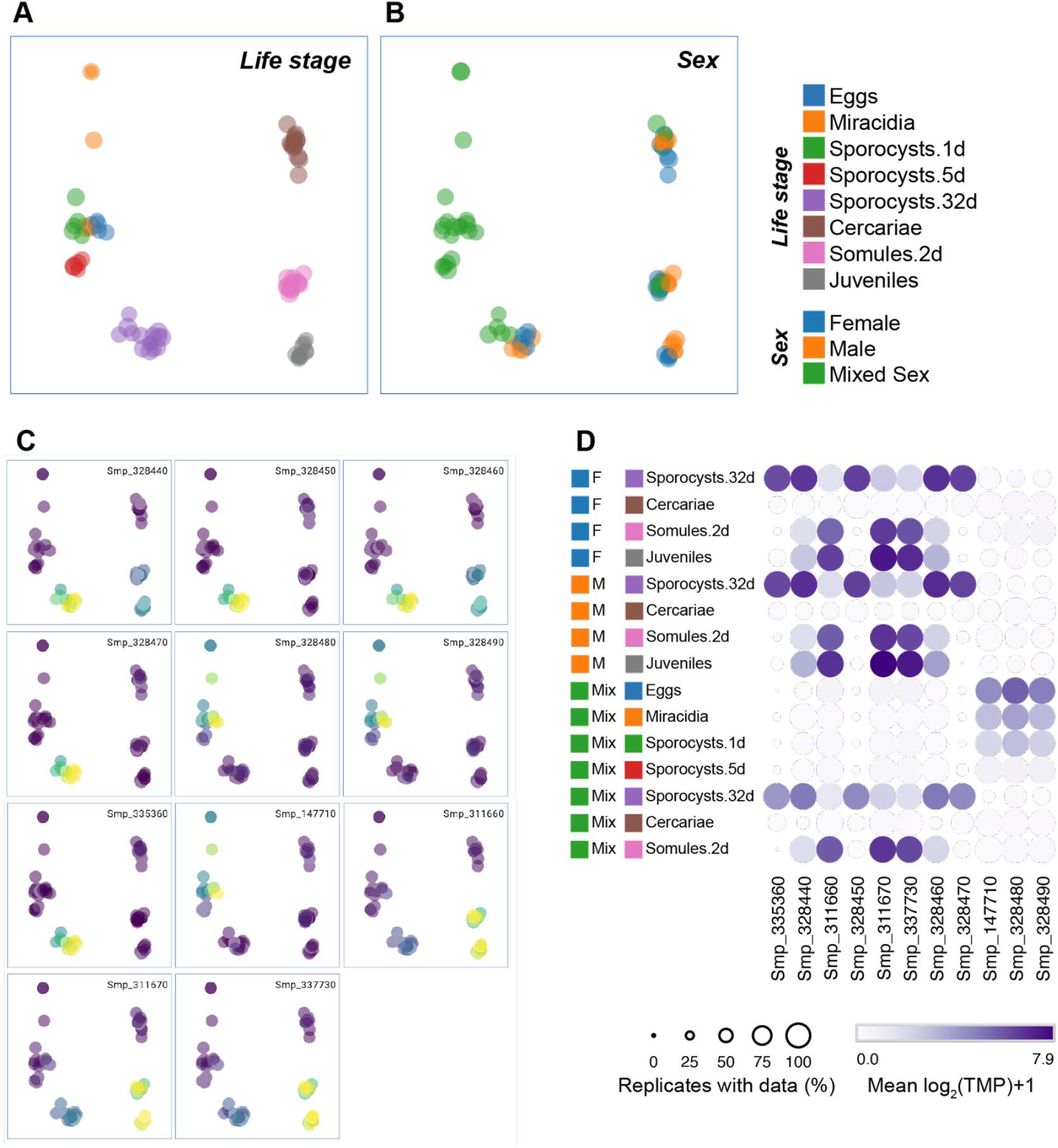
Visualisation and exploration of stage and sex-specific RNA-seq data of *Schistosoma mansoni*. Demonstration of our easy-to-use data exploration tool https://lifecycle.schisto.xyz/ using *S. mansoni* Kunitz protease inhibitors as an example. **(A, B)** PCA plot of RNA-seq data coloured by life stage and by parasite sex, highlighting male, female, and mixe-sex parasites, to provide context for gene-specific analyses. **(C)** Series of PCA plots showing the expression of 11 *S. mansoni* Kunitz protease inhibitors. In A, B, and C, each point represents a single replicate (n = 75) and is coloured by log transcript per million values (log_2_[TPM+1]), with low expression represented by dark and high expression light colour. **(D)** A Heatmap of mean log_2_(TPM+1) expression for each gene is presented in panel C. Each point represents the mean TMP from five replicates, with the size of the point proportional to the number of replicates within a group with data.

## Code Availability

All code used to analyse this data set has been deposited in the GitHub repository https://github.com/SKBuddenborg/Smansoni_rnaseq and is archived at Zenodo (DOI 10.5281/zenodo.7849283)^40^.

## Acknowledgements

We thank the Pathogen Informatics group at the Wellcome Sanger Institute for informatics support. We also thank Gabriel Rinaldi and other members of the Berriman Lab for discussions and support with the maintenance of the *S. mansoni* life cycle. This work was supported by the Wellcome Trust through core funding to the Wellcome Sanger Institute [206194]. SRD is supported by a UKRI Future Leaders Fellowship [MR/T020733/1]. For the purpose of Open Access, the author has applied a CC BY public copyright licence to any Author Accepted Manuscript version arising from this submission.

## Author contributions

SKB and MB conceived and designed the study, SKB and GS performed the experiments, SKB analysed RNA-seq data, ZL supported the data visualisation, SRD supervised data analysis and wrote the manuscript with SKB, with contributions from the other authors.

