## Supplemental Table 1 for "The stage- and sex-specific transcriptome of the human parasite *Schistosoma mansoni*"

| Internal Sample ID | Library Name | Sample Name | ENA Sample Accession | i7 Tag Sequence | i5 Tag Sequence | Lane 1 filename | Lane 1 ENA Run Accession | Lane 1 read count | Total filtered reads | Lane 1 uniquely mapped | Lane 2 filename | Lane 2 ENA Run Accession | Lane 2 Read count | Total filtered reads | Lane 2 unique mapped | Total read count | Total v10 PE mapped reads | Percent v10 PE mapped reads | Location | Sex |
|---|---|---|---|---|---|---|---|---|---|---|---|---|---|---|---|---|---|---|---|---|
| 5837STDY9141995 | NT1631279D | X_Eggs_R1 | ERS4985650 | TTACCGAC | CGAATACG | 35101_2#3 | ERR11178266 | 35,887,713 | 35,711,893 | 28,792,320 | 35207_2#3 | ERR11178356 | 37,143,727 | 34,707,754 | 28,008,863 | 73,031,440 | 56,801,183 | 77.78 | Intramammalian | Mix |
| 5837STDY9141996 | NT1631280T | X_Eggs_R2 | ERS4985649 | TCGTCTGA | GTCCTTGA | 35101_2#13 | ERR11178276 | 29,267,011 | 27,926,030 | 23,410,952 | 35207_2#13 | ERR11178366 | 30,482,693 | 27,197,392 | 22,836,299 | 59,749,704 | 46,247,251 | 77.40 | Intramammalian | Mix |
| 5837STDY9141997 | NT1631281U | X_Eggs_R3 | ERS4985652 | TTCCAGGT | CAGTGCTT | 35101_2#23 | ERR11178286 | 38,431,463 | 38,662,920 | 31,139,740 | 35207_2#23 | ERR11178376 | 39,962,345 | 37,370,199 | 30,121,675 | 78,393,808 | 61,261,415 | 78.15 | Intramammalian | Mix |
| 5837STDY9141998 | NT1631282V | X_Eggs_R4 | ERS4985654 | TACGGTCT | TCCATTGC | 35101_2#32 | ERR11178295 | 38,386,940 | 37,603,277 | 31,092,926 | 35207_2#32 | ERR11178385 | 40,013,153 | 36,421,944 | 30,118,014 | 78,400,093 | 61,210,940 | 78.08 | Intramammalian | Mix |
| 5837STDY9141999 | NT1631283W | X_Eggs_R5 | ERS4985656 | AAGACCGT | GTCGATTG | 35101_2#41 | ERR11178304 | 33,520,934 | 33,370,668 | 27,676,614 | 35207_2#41 | ERR11178394 | 34,292,942 | 32,800,293 | 27,223,668 | 67,813,876 | 54,900,282 | 80.96 | Intramammalian | Mix |
| 5837STDY9142000 | NT1631284A | X_Miracidia_R1 | ERS4985658 | CAGGTTCA | ATAACGCC | 35101_2#50 | ERR11178313 | 11,810,920 | 11,081,039 | 9,238,789 | 35207_2#50 | ERR11178403 | 12,515,180 | 10,607,443 | 8,857,137 | 24,326,100 | 18,095,926 | 74.39 | Free-living, water | Mix |
| 5837STDY9142001 | NT1631285B | X_Miracidia_R2 | ERS4985660 | TAGGAGCT | GCCTTAAC | 35101_2#59 | ERR11178322 | 91,165,702 | 89,868,791 | 77,502,575 | 35207_2#59 | ERR11178412 | 92,175,872 | 89,282,000 | 77,003,851 | 183,341,574 | 154,506,426 | 84.27 | Free-living, water | Mix |
| 5837STDY9142002 | NT1631286C | X_Miracidia_R3 | ERS4985661 | TACTCCAG | GGTATAGG | 35101_2#67 | ERR11178330 | 25,106,884 | 12,022,876 | 9,367,104 | 35207_2#67 | ERR11178420 | 28,922,717 | 11,428,849 | 8,972,765 | 54,029,601 | 18,339,869 | 33.94 | Free-living, water | Mix |
| 5837STDY9142003 | NT1631287D | X_Miracidia_R4 | ERS4985664 | AGTGACCT | TCTAGGAG | 35101_2#5 | ERR11178268 | 31,286,811 | 31,008,863 | 26,656,759 | 35207_2#5 | ERR11178358 | 32,146,173 | 30,361,820 | 26,117,974 | 63,432,984 | 52,774,733 | 83.20 | Free-living, water | Mix |
| 5837STDY9142004 | NT1631288E | X_Miracidia_R5 | ERS4985665 | AGCCTATC | TGCGTAAC | 35101_2#15 | ERR11178278 | 36,697,581 | 36,876,449 | 31,613,336 | 35207_2#15 | ERR11178368 | 38,496,384 | 35,399,825 | 30,362,483 | 75,193,965 | 61,975,819 | 82.42 | Free-living, water | Mix |
| 5837STDY9142005 | NT1631289F | X_1d_Sporocysts_R1 | ERS4985668 | TCATCTCC | CTTGCTAG | 35101_2#25 | ERR11178288 | 32,277,847 | 31,838,943 | 26,172,977 | 35207_2#25 | ERR11178378 | 33,963,863 | 30,541,659 | 25,135,832 | 66,241,710 | 51,308,809 | 77.46 | *in vitro* | Mix |
| 5837STDY9142006 | NT1631290V | X_1d_Sporocysts_R2 | ERS4985670 | CCAGTATC | AGCGAGAT | 35101_2#34 | ERR11178297 | 35,892,793 | 35,677,673 | 29,621,370 | 35207_2#34 | ERR11178387 | 37,594,051 | 34,325,361 | 28,519,219 | 73,486,844 | 58,140,589 | 79.12 | *in vitro* | Mix |
| 5837STDY9142007 | NT1631291W | X_1d_Sporocysts_R3 | ERS4985672 | TTGCGAGA | TATGGCAC | 35101_2#43 | ERR11178306 | 35,017,895 | 35,134,512 | 28,942,807 | 35207_2#43 | ERR11178396 | 36,817,188 | 33,654,597 | 27,742,076 | 71,835,083 | 56,684,883 | 78.91 | *in vitro* | Mix |
| 5837STDY9142008 | NT1631292A | X_1d_Sporocysts_R4 | ERS4985674 | GAACGAAG | GAATCACC | 35101_2#52 | ERR11178315 | 29,641,288 | 29,545,049 | 24,067,975 | 35207_2#52 | ERR11178405 | 30,864,285 | 28,558,053 | 23,275,383 | 60,505,573 | 47,343,358 | 78.25 | *in vitro* | Mix |
| 5837STDY9142009 | NT1631293B | X_1d_Sporocysts_R5 | ERS4985676 | CGAATTGC | GTAAGGTG | 35101_2#1 | ERR11178264 | 30,832,268 | 31,868,421 | 26,041,582 | 35207_2#1 | ERR11178354 | 33,177,814 | 29,848,971 | 24,422,313 | 64,010,082 | 50,463,895 | 78.84 | *in vitro* | Mix |
| 5837STDY9142010 | NT1631294C | X_5d_Sporocysts_R1 | ERS4985677 | GGAAGAGA | CGAGAGAA | 35101_2#69 | ERR11178332 | 27,466,726 | 27,362,950 | 22,761,139 | 35207_2#69 | ERR11178422 | 28,178,124 | 26,804,403 | 22,302,697 | 55,644,850 | 45,063,836 | 80.98 | *in vitro* | Mix |
| 5837STDY9142011 | NT1631295D | X_5d_Sporocysts_R2 | ERS4985679 | TCGGATTC | CGCAACTA | 35101_2#7 | ERR11178270 | 28,967,618 | 27,007,667 | 22,216,700 | 35207_2#7 | ERR11178360 | 30,375,494 | 26,234,530 | 21,609,581 | 59,343,112 | 43,826,281 | 73.85 | *in vitro* | Mix |
| 5837STDY9142012 | NT1631296E | X_5d_Sporocysts_R3 | ERS4985680 | CTGTACCA | CACAGACT | 35101_2#17 | ERR11178280 | 35,109,519 | 33,827,742 | 28,333,428 | 35207_2#17 | ERR11178370 | 35,920,899 | 33,385,587 | 27,971,592 | 71,030,418 | 56,305,020 | 79.27 | *in vitro* | Mix |
| 5837STDY9142013 | NT1631297F | X_5d_Sporocysts_R4 | ERS4985683 | GAGAGTAC | TGGAAGCA | 35101_2#27 | ERR11178290 | 36,963,589 | 36,157,533 | 30,869,844 | 35207_2#27 | ERR11178380 | 38,214,917 | 35,270,776 | 30,123,650 | 75,178,506 | 60,993,494 | 81.13 | *in vitro* | Mix |
| 5837STDY9142014 | NT1631298G | X_5d_Sporocysts_R5 | ERS4985685 | TCTACGCA | CAATAGCC | 35101_2#36 | ERR11178299 | 37,279,602 | 36,223,301 | 30,582,388 | 35207_2#36 | ERR11178389 | 37,972,790 | 35,830,165 | 30,252,589 | 75,252,392 | 60,834,977 | 80.84 | *in vitro* | Mix |
| 5837STDY9142015 | NT1631299H | F_32d_Sporocysts_R1 | ERS4985687 | GCAATTCC | CTCGAACA | 35101_2#45 | ERR11178308 | 19,916,240 | 19,860,827 | 4,503,158 | 35207_2#45 | ERR11178398 | 20,647,759 | 19,277,995 | 4,383,669 | 40,563,999 | 8,886,827 | 21.91 | Intramolluscan | Female |
| 5837STDY9142016 | NT1631300G | F_32d_Sporocysts_R2 | ERS4985689 | CTCAGAAG | GGCAAGTT | 35101_2#54 | ERR11178317 | 23,616,338 | 23,535,954 | 6,501,038 | 35207_2#54 | ERR11178407 | 24,220,564 | 23,050,267 | 6,385,220 | 47,836,902 | 12,886,258 | 26.94 | Intramolluscan | Female |
| 5837STDY9142017 | NT1631301H | F_32d_Sporocysts_R3 | ERS4985691 | GTCCTAAG | AGCTACCA | 35101_2#62 | ERR11178325 | 32,154,542 | 31,690,547 | 8,172,841 | 35207_2#62 | ERR11178415 | 33,641,955 | 30,572,065 | 7,905,397 | 65,796,497 | 16,078,238 | 24.44 | Intramolluscan | Female |
| 5837STDY9142018 | NT1631302I | F_32d_Sporocysts_R4 | ERS4985693 | GCGTTAGA | CAGCATAC | 35101_2#71 | ERR11178334 | 29,669,554 | 27,851,289 | 7,442,478 | 35207_2#71 | ERR11178424 | 31,162,150 | 27,008,423 | 7,238,358 | 60,831,704 | 14,680,836 | 24.13 | Intramolluscan | Female |
| 5837STDY9142019 | NT1631303J | F_32d_Sporocysts_R5 | ERS4985695 | CAAGGTAC | CGTATCTC | 35101_2#9 | ERR11178272 | 25,041,123 | 24,578,344 | 8,193,221 | 35207_2#9 | ERR11178362 | 25,491,812 | 24,299,568 | 8,119,470 | 50,532,935 | 16,312,691 | 32.28 | Intramolluscan | Female |
| 5837STDY9142020 | NT1631304K | M_32d_Sporocysts_R1 | ERS4985697 | AGACCTTG | TTACGTGC | 35101_2#19 | ERR11178282 | 20,507,747 | 21,046,824 | 5,627,026 | 35207_2#19 | ERR11178372 | 21,459,171 | 20,172,481 | 5,416,731 | 41,966,918 | 11,043,757 | 26.32 | Intramolluscan | Male |
| 5837STDY9142021 | NT1631305L | M_32d_Sporocysts_R2 | ERS4985699 | GTCGTTAC | AGCTAAGC | 35101_2#29 | ERR11178292 | 33,594,820 | 31,819,567 | 8,954,939 | 35207_2#29 | ERR11178382 | 35,115,864 | 30,912,846 | 8,705,919 | 68,710,684 | 17,660,858 | 25.70 | Intramolluscan | Male |
| 5837STDY9142022 | NT1631306M | M_32d_Sporocysts_R3 | ERS4985701 | GTAACCGA | AAGACACC | 35101_2#38 | ERR11178301 | 24,052,782 | 24,774,896 | 6,701,453 | 35207_2#38 | ERR11178391 | 25,144,038 | 23,743,243 | 6,436,225 | 49,196,820 | 13,137,678 | 26.70 | Intramolluscan | Male |
| 5837STDY9142023 | NT1631307N | M_32d_Sporocysts_R4 | ERS4985704 | GAATCCGT | CAACTCCA | 35101_2#47 | ERR11178310 | 52,577,317 | 52,982,669 | 9,133,060 | 35207_2#47 | ERR11178400 | 53,752,992 | 51,944,602 | 8,963,302 | 106,330,309 | 18,096,362 | 17.02 | Intramolluscan | Male |
| 5837STDY9142024 | NT1631308O | M_32d_Sporocysts_R5 | ERS4985705 | CATGAGCA | GATCTTGC | 35101_2#56 | ERR11178319 | 33,075,456 | 32,849,017 | 11,294,032 | 35207_2#56 | ERR11178409 | 34,384,073 | 31,799,664 | 10,955,823 | 67,459,529 | 22,249,855 | 32.98 | Intramolluscan | Male |
| 5837STDY9142025 | NT1631309P | X_32d_Sporocysts_R1 | ERS4985707 | CTTAGGAC | CTTCACTG | 35101_2#64 | ERR11178327 | 52,413,949 | 53,349,468 | 17,985,300 | 35207_2#64 | ERR11178417 | 54,246,010 | 51,686,663 | 17,438,802 | 106,659,959 | 35,424,102 | 33.21 | Intramolluscan | Mix |
| 5837STDY9142026 | NT1631310I | X_32d_Sporocysts_R2 | ERS4985708 | ATCTGACC | CTCGACTT | 35101_2#73 | ERR11178336 | 27,571,604 | 27,328,402 | 10,035,045 | 35207_2#73 | ERR11178426 | 28,284,142 | 26,780,391 | 9,855,024 | 55,855,746 | 19,890,069 | 35.61 | Intramolluscan | Mix |
| 5837STDY9142027 | NT1631311J | X_32d_Sporocysts_R3 | ERS4985711 | TCCTCATG | GTACACCT | 35101_2#78 | ERR11178341 | 21,105,095 | 20,382,083 | 5,522,652 | 35207_2#78 | ERR11178431 | 22,167,144 | 19,649,735 | 5,330,092 | 43,272,239 | 10,852,744 | 25.08 | Intramolluscan | Mix |
| 5837STDY9142028 | NT1631312K | X_32d_Sporocysts_R4 | ERS4985713 | AGGATAGC | CCAAGGTT | 35101_2#82 | ERR11178345 | 19,987,672 | 21,643,912 | 6,507,569 | 35207_2#82 | ERR11178435 | 20,797,374 | 20,685,494 | 6,228,954 | 40,785,046 | 12,736,523 | 31.23 | Intramolluscan | Mix |
| 5837STDY9142029 | NT1631313L | X_32d_Sporocysts_R5 | ERS4985715 | GGAGGAAT | GAACGGTT | 35101_2#79 | ERR11178342 | 12,935,454 | 13,175,297 | 5,007,826 | 35207_2#79 | ERR11178432 | 13,397,086 | 12,756,662 | 4,845,642 | 26,332,540 | 9,853,468 | 37.42 | Intramolluscan | Mix |
| 5837STDY9142030 | NT1631314M | F_Cercariae_R1 | ERS4985717 | GACGTCAT | CCAGTTGA | 35101_2#80 | ERR11178343 | 15,787,887 | 14,160,908 | 12,052,642 | 35207_2#80 | ERR11178433 | 16,220,056 | 14,087,751 | 12,016,455 | 32,007,943 | 24,069,097 | 75.20 | Free-living, water | Female |
| 5837STDY9142031 | NT1631315N | F_Cercariae_R2 | ERS4985719 | CCGCTTAA | GTCATCGT | 35101_2#84 | ERR11178347 | 17,756,605 | 17,194,805 | 13,946,659 | 35207_2#84 | ERR11178437 | 18,403,434 | 16,757,409 | 13,622,626 | 36,160,039 | 27,569,285 | 76.24 | Free-living, water | Female |
| 5837STDY9142032 | NT1631316O | F_Cercariae_R3 | ERS4985722 | GACGAACT | CAATGCGA | 35101_2#86 | ERR11178349 | 15,360,630 | 14,867,063 | 12,388,092 | 35207_2#86 | ERR11178439 | 15,825,906 | 14,567,299 | 12,153,590 | 31,186,536 | 24,541,682 | 78.69 | Free-living, water | Female |
| 5837STDY9142033 | NT1631317P | F_Cercariae_R4 | ERS4985723 | TCCACGTT | GGTTGAAC | 35101_2#88 | ERR11178351 | 18,223,529 | 16,946,350 | 14,034,844 | 35207_2#88 | ERR11178441 | 18,644,522 | 16,839,984 | 13,974,698 | 36,868,051 | 28,009,542 | 75.97 | Free-living, water | Female |
| 5837STDY9142034 | NT1631318Q | F_Cercariae_R5 | ERS4985727 | AACCAGAG | CTTCGGTT | 35101_2#90 | ERR11178353 | 22,046,492 | 20,009,204 | 16,083,050 | 35207_2#90 | ERR11178443 | 22,853,131 | 19,679,706 | 15,856,698 | 44,899,623 | 31,939,748 | 71.14 | Free-living, water | Female |
| 5837STDY9142035 | NT1631319R | M_Cercariae_R1 | ERS4985729 | GTCAGTCA | CGGCATTA | 35101_2#11 | ERR11178274 | 15,123,476 | 12,621,823 | 10,481,420 | 35207_2#11 | ERR11178364 | 16,147,537 | 12,189,372 | 10,147,144 | 31,271,013 | 20,628,564 | 65.97 | Free-living, water | Male |
| 5837STDY9142036 | NT1631320K | M_Cercariae_R2 | ERS4985731 | CCTTCCAT | CACGCAAT | 35101_2#21 | ERR11178284 | 22,997,858 | 19,473,971 | 16,080,101 | 35207_2#21 | ERR11178374 | 24,418,934 | 19,008,147 | 15,727,336 | 47,416,792 | 31,807,437 | 67.08 | Free-living, water | Male |
| 5837STDY9142037 | NT1631321L | M_Cercariae_R3 | ERS4985733 | AGGAACAC | GGAATGTC | 35101_2#30 | ERR11178293 | 22,111,013 | 21,163,731 | 17,881,725 | 35207_2#30 | ERR11178383 | 23,083,597 | 20,543,863 | 17,381,539 | 45,194,610 | 35,263,264 | 78.03 | Free-living, water | Male |
| 5837STDY9142038 | NT1631322M | M_Cercariae_R4 | ERS4985735 | CTTACAGC | TGGTGAAG | 35101_2#39 | ERR11178302 | 27,749,338 | 25,192,733 | 20,992,854 | 35207_2#39 | ERR11178392 | 29,793,050 | 24,038,611 | 20,064,889 | 57,542,388 | 41,057,743 | 71.35 | Free-living, water | Male |
| 5837STDY9142039 | NT1631323N | M_Cercariae_R5 | ERS4985737 | TACCTGCA | GGACATCA | 35101_2#48 | ERR11178311 | 24,028,945 | 23,497,244 | 19,865,693 | 35207_2#48 | ERR11178401 | 25,097,280 | 22,711,959 | 19,227,421 | 49,126,225 | 39,093,114 | 79.58 | Free-living, water | Male |
| 5837STDY9142040 | NT1631324O | X_Cercariae_R1 | ERS4985739 | AGACGCTA | GGTGTACA | 35101_2#57 | ERR11178320 | 30,839,479 | 29,309,804 | 22,920,893 | 35207_2#57 | ERR11178410 | 31,205,940 | 29,140,689 | 22,922,607 | 62,045,419 | 45,843,500 | 73.89 | Free-living, water | Mix |
| 5837STDY9142041 | NT1631325P | X_Cercariae_R2 | ERS4985741 | CAACACAG | GATAGCCA | 35101_2#65 | ERR11178328 | 21,447,923 | 20,400,740 | 16,003,015 | 35207_2#65 | ERR11178418 | 22,460,049 | 19,771,388 | 15,520,605 | 43,907,972 | 31,523,620 | 71.79 | Free-living, water | Mix |
| 5837STDY9142042 | NT1631326Q | X_Cercariae_R3 | ERS4985725 | GTACCACA | CCACAACA | 35101_2#74 | ERR11178337 | 25,576,960 | 24,825,597 | 19,749,246 | 35207_2#74 | ERR11178427 | 26,420,267 | 24,259,936 | 19,305,925 | 51,997,227 | 39,055,171 | 75.11 | Free-living, water | Mix |
| 5837STDY9142043 | NT1631327R | X_Cercariae_R4 | ERS4985726 | CGAATACG | TTACCGAC | 35101_2#2 | ERR11178265 | 25,094,629 | 22,206,698 | 16,561,221 | 35207_2#2 | ERR11178355 | 27,381,310 | 20,917,747 | 15,620,455 | 52,475,939 | 32,181,676 | 61.33 | Free-living, water | Mix |
| 5837STDY9142044 | NT1631328S | X_Cercariae_R5 | ERS4985743 | GTCCTTGA | TCGTCTGA | 35101_2#12 | ERR11178275 | 27,558,724 | 25,708,095 | 19,071,232 | 35207_2#12 | ERR11178365 | 29,228,943 | 24,682,820 | 18,318,320 | 56,787,667 | 37,389,552 | 65.84 | Free-living, water | Mix |
| 5837STDY9142045 | NT1631329T | F_2d_Somules_R1 | ERS4985745 | CAGTGCTT | TTCCAGGT | 35101_2#22 | ERR11178285 | 37,217,104 | 30,087,711 | 26,168,037 | 35207_2#22 | ERR11178375 | 39,335,841 | 29,643,158 | 25,832,243 | 76,552,945 | 52,000,280 | 67.93 | *in vitro* | Female |
| 5837STDY9142046 | NT1631330M | F_2d_Somules_R2 | ERS4985747 | TCCATTGC | TACGGTCT | 35101_2#31 | ERR11178294 | 21,219,579 | 19,906,136 | 14,763,901 | 35207_2#31 | ERR11178384 | 22,710,301 | 18,952,426 | 14,069,976 | 43,929,880 | 28,833,877 | 65.64 | *in vitro* | Female |
| 5837STDY9142047 | NT1631331N | F_2d_Somules_R3 | ERS4985749 | GTCGATTG | AAGACCGT | 35101_2#40 | ERR11178303 | 21,234,685 | 19,417,533 | 16,365,076 | 35207_2#40 | ERR11178393 | 23,097,477 | 18,263,122 | 15,404,488 | 44,332,162 | 31,769,564 | 71.66 | *in vitro* | Female |
| 5837STDY9142048 | NT1631332O | F_2d_Somules_R4 | ERS4985751 | ATAACGCC | CAGGTTCA | 35101_2#49 | ERR11178312 | 33,135,155 | 23,844,782 | 19,823,869 | 35207_2#49 | ERR11178402 | 35,463,886 | 23,361,646 | 19,460,453 | 68,599,041 | 39,284,322 | 57.27 | *in vitro* | Female |
| 5837STDY9142049 | NT1631333P | F_2d_Somules_R5 | ERS4985753 | GCCTTAAC | TAGGAGCT | 35101_2#58 | ERR11178321 | 26,946,916 | 21,935,535 | 17,606,289 | 35207_2#58 | ERR11178411 | 29,777,996 | 20,680,360 | 16,623,469 | 56,724,912 | 34,229,758 | 60.34 | *in vitro* | Female |
| 5837STDY9142050 | NT1631334Q | M_2d_Somules_R1 | ERS4985755 | GGTATAGG | TACTCCAG | 35101_2#66 | ERR11178329 | 47,675,928 | 34,126,982 | 30,187,161 | 35207_2#66 | ERR11178419 | 52,708,438 | 32,616,311 | 28,940,980 | 100,384,366 | 59,128,141 | 58.90 | *in vitro* | Male |
| 5837STDY9142051 | NT1631335R | M_2d_Somules_R2 | ERS4985757 | TCTAGGAG | AGTGACCT | 35101_2#4 | ERR11178267 | 29,742,096 | 29,713,378 | 26,920,258 | 35207_2#4 | ERR11178357 | 31,501,829 | 28,306,929 | 25,657,000 | 61,243,925 | 52,577,258 | 85.85 | *in vitro* | Male |

| | | | | | | | | | | | | | | | | | | | | |
|---|---|---|---|---|---|---|---|---|---|---|---|---|---|---|---|---|---|---|---|---|
| 5837STDY9142052 | NT1631336S | M_2d_Somules_R3 | ERS4985759 | TGCGTAAC | AGCCTATC | 35101_2#14 | ERR11178277 | 24,598,676 | 24,130,291 | 21,661,079 | 35207_2#14 | ERR11178367 | 25,978,279 | 23,116,192 | 20,760,829 | 50,576,955 | 42,421,908 | 83.88 | *in vitro* | Male |
| 5837STDY9142053 | NT1631337T | M_2d_Somules_R4 | ERS4985761 | CTTGCTAG | TCATCTCC | 35101_2#24 | ERR11178287 | 14,446,390 | 14,404,395 | 12,880,823 | 35207_2#24 | ERR11178377 | 15,329,895 | 13,694,470 | 12,241,807 | 29,776,285 | 25,122,630 | 84.37 | *in vitro* | Male |
| 5837STDY9142054 | NT1631338U | M_2d_Somules_R5 | ERS4985763 | AGCGAGAT | CCAGTATC | 35101_2#33 | ERR11178296 | 23,623,195 | 22,703,108 | 20,432,867 | 35207_2#33 | ERR11178386 | 24,093,106 | 22,478,132 | 20,229,990 | 47,716,301 | 40,662,857 | 85.22 | *in vitro* | Male |
| 5837STDY9142055 | NT1631339V | X_2d_Somules_R1 | ERS4985765 | TATGGCAC | TTGCGAGA | 35101_2#42 | ERR11178305 | 39,081,876 | 37,554,727 | 33,344,456 | 35207_2#42 | ERR11178395 | 40,661,718 | 36,502,728 | 32,422,150 | 79,743,594 | 65,766,606 | 82.47 | *in vitro* | Mix |
| 5837STDY9142056 | NT1631340O | X_2d_Somules_R2 | ERS4985766 | GAATCACC | GAACGAAG | 35101_2#51 | ERR11178314 | 41,779,703 | 41,767,124 | 37,118,495 | 35207_2#51 | ERR11178404 | 43,666,563 | 40,249,600 | 35,779,082 | 85,446,266 | 72,897,577 | 85.31 | *in vitro* | Mix |
| 5837STDY9142057 | NT1631341P | X_2d_Somules_R3 | ERS4985770 | GTAAGGTG | CGAATTGC | 35101_2#60 | ERR11178323 | 56,749,551 | 58,027,334 | 52,046,890 | 35207_2#60 | ERR11178413 | 60,015,644 | 55,140,052 | 49,471,995 | 116,765,195 | 101,518,885 | 86.94 | *in vitro* | Mix |
| 5837STDY9142068 | NT1631352S | X_2d_Somules_R4 | ERS4985787 | TTACGTGC | AGACCTTG | 35101_2#18 | ERR11178281 | 22,305,485 | 22,113,245 | 19,598,230 | 35207_2#18 | ERR11178371 | 23,607,071 | 21,119,969 | 18,717,132 | 45,912,556 | 38,315,362 | 83.45 | *in vitro* | Mix |
| 5837STDY9142069 | NT1631353T | X_2d_Somules_R5 | ERS4985788 | AGCTAAGC | GTCGTTAC | 35101_2#28 | ERR11178291 | 14,700,276 | 14,402,717 | 12,629,923 | 35207_2#28 | ERR11178381 | 15,177,272 | 14,068,732 | 12,333,592 | 29,877,548 | 24,963,515 | 83.55 | *in vitro* | Mix |
| 5837STDY9142058 | NT1631342Q | F_26d_Juveniles_R1 | ERS4985792 | CGAGAGAA | GGAAGAGA | 35101_2#68 | ERR11178331 | 37,617,772 | 37,486,481 | 33,544,742 | 35207_2#68 | ERR11178421 | 39,296,367 | 36,126,210 | 32,334,070 | 76,914,139 | 65,878,812 | 85.65 | Intramammalian | Female |
| 5837STDY9142059 | NT1631343R | F_26d_Juveniles_R2 | ERS4985793 | CGCAACTA | TCGGATTC | 35101_2#6 | ERR11178269 | 18,817,707 | 19,037,704 | 17,118,565 | 35207_2#6 | ERR11178359 | 19,666,730 | 18,306,076 | 16,459,770 | 38,484,437 | 33,578,335 | 87.25 | Intramammalian | Female |
| 5837STDY9142060 | NT1631344S | F_26d_Juveniles_R3 | ERS4985771 | CACAGACT | CTGTACCA | 35101_2#16 | ERR11178279 | 16,570,080 | 16,304,076 | 14,610,323 | 35207_2#16 | ERR11178369 | 17,259,493 | 15,787,215 | 14,151,732 | 33,829,573 | 28,762,055 | 85.02 | Intramammalian | Female |
| 5837STDY9142061 | NT1631345T | F_26d_Juveniles_R4 | ERS4985773 | TGGAAGCA | GAGAGTAC | 35101_2#26 | ERR11178289 | 10,488,945 | 10,426,601 | 9,353,583 | 35207_2#26 | ERR11178379 | 10,826,459 | 10,158,814 | 9,118,832 | 21,315,404 | 18,472,415 | 86.66 | Intramammalian | Female |
| 5837STDY9142062 | NT1631346U | F_26d_Juveniles_R5 | ERS4985776 | CAATAGCC | TCTACGCA | 35101_2#35 | ERR11178298 | 32,411,201 | 32,308,290 | 29,005,197 | 35207_2#35 | ERR11178388 | 33,549,366 | 31,405,770 | 28,198,400 | 65,960,567 | 57,203,597 | 86.72 | Intramammalian | Female |
| 5837STDY9142063 | NT1631347V | M_26d_Juveniles_R1 | ERS4985777 | CTCGAACA | GCAATTCC | 35101_2#44 | ERR11178307 | 13,434,861 | 13,722,260 | 12,312,366 | 35207_2#44 | ERR11178397 | 13,998,964 | 13,211,220 | 11,852,416 | 27,433,825 | 24,164,782 | 88.08 | Intramammalian | Male |
| 5837STDY9142064 | NT1631348W | M_26d_Juveniles_R2 | ERS4985779 | GGCAAGTT | CTCAGAAG | 35101_2#53 | ERR11178316 | 20,880,475 | 20,271,512 | 18,069,197 | 35207_2#53 | ERR11178406 | 20,871,333 | 20,377,326 | 18,170,472 | 41,751,808 | 36,239,669 | 86.80 | Intramammalian | Male |
| 5837STDY9142065 | NT1631349A | M_26d_Juveniles_R3 | ERS4985780 | AGCTACCA | GTCCTAAG | 35101_2#61 | ERR11178324 | 33,691,182 | 32,928,974 | 29,345,165 | 35207_2#61 | ERR11178414 | 34,135,123 | 32,699,524 | 29,154,608 | 67,826,305 | 58,499,773 | 86.25 | Intramammalian | Male |
| 5837STDY9142066 | NT1631350Q | M_26d_Juveniles_R4 | ERS4985784 | CAGCATAC | GCGTTAGA | 35101_2#70 | ERR11178333 | 29,106,042 | 28,585,962 | 25,552,867 | 35207_2#70 | ERR11178423 | 29,375,432 | 28,451,574 | 25,429,012 | 58,481,474 | 50,981,879 | 87.18 | Intramammalian | Male |
| 5837STDY9142067 | NT1631351R | M_26d_Juveniles_R5 | ERS4985785 | CGTATCTC | CAAGGTAC | 35101_2#8 | ERR11178271 | 31,868,656 | 32,102,065 | 28,643,522 | 35207_2#8 | ERR11178361 | 33,505,258 | 30,760,007 | 27,458,468 | 65,373,914 | 56,101,990 | 85.82 | Intramammalian | Male |
